# *Salmonella* grows massively and aerobically in fecal matter

**DOI:** 10.1101/766782

**Authors:** Teresa Guerrero, Sonia Zapata, Gabriel Trueba

**Affiliations:** Microbiology Institute, Universidad San Francisco de Quito, Diego de Robles y Vía Interoceánica, Cumbayá, Quito, Ecuador

## Abstract

The use of wastewater for irrigation and animal manure as fertilizer can cause transmission of intestinal pathogens, conditions frequently observed in Low- and Middle-Income Countries (LMICs). Here we tested the ability of *Salmonella* to grow in the fecal matter; we inoculated freshly isolated *Salmonella* strains (from chickens) in chicken fecal matter and incubated for 24, 48 and 72 hrs under aerobic and anaerobic conditions. We found that both *Salmonella* and *E. coli* multiplied massively in fecal matter outside a host for 72 hrs, being their growth higher in aerobic conditions. Our results have critical implications in waste management, as we demonstrate that aerobic treatments may not be the best to reduce the number of *Salmonella* in the environment.

## Introduction

Environmental transmission of intestinal pathogens is extremely important especially in Low- and Middle-Income Countries (LMICs) due to deficient sanitary infrastructure, unplanned urban growth, lack of wastewater treatment, etc. One of the main concerns in LMICs is the large proportion of untreated wastewater used for irrigation (Khalid *et al*., 2018) and the increasing use of animal manure as fertilizer without suitable treatment (Mandrell, 2009); these are problems that remain neglected in LMICs (Khalid *et al*., 2018). Reports of grave enteric infections caused by environmental contamination of produce are also commonplace nowadays in industrialized countries (Callejón *et al*., 2015). Some of these outbreaks have been associated with high mortality, morbidity and large economic losses (Mandrell, 2009). The incidence of these infections is exacerbated by the increasing appeal to consume natural, non-processed fresh products (Mandrell, 2009).

*Salmonella* contaminated water is responsible for a large number of outbreaks by ingestion of water or produce (Mandrell, 2009); the sources for this contamination are human and non-human fecal matter (Medrano-Félix *et al*., 2017). The use of animal waste as fertilizer constitutes a serious risk which can be controlled by appropriate composting technology (Tiquia *et al*., 1998; Szogi *et al.*, 2015). Human waste contamination, however, is much more difficult to monitor or control in LMICs where wastewater treatment or toilets are not available (Khalid *et al*., 2018); the fate of *Salmonella* in these conditions is not understood completely, although some researchers indicate that *Salmonella* enters into a viable-non culturable state outside the host (Winfield and Groisman, 2003). The reduction of the risk of this type of transmission requires the understanding of every aspect of *Salmonella* physiology in the environment outside the host (Mandrell, 2009). It is worth mentioning that *Salmonella*’s ability to grow in fecal matter has been ignored.

It is known that *Salmonella* and other *Enterobacteriaceae* survive in a fecal matter for some time and it has been shown that *E. coli* (another member of the *Enterobacteriaceae*) also grows massively in fecal matter (Russell and Jarvis, 2001; Vasco *et al.*, 2015; Sharma *et al*., 2019). Here we tested *Salmonella*’s ability to grow in fecal matter in aerobic conditions and discuss the potential implications for fecal waste management.

## Experimental Procedures

### Overall approach

We inoculated fresh *Salmonella* isolates in chicken fecal matter and incubated for 24, 48 and 72 hrs under aerobic and anaerobic conditions; we counted *Salmonella* colonies number by culture and by the loop mediated isothermal amplification 3M™ Molecular Detection Assay 2 - Salmonella (MDA2SAL).

Five fresh *Salmonella* isolates, obtained from poultry, were identified as *S. enterica* serovars Infantis, Dublin, Heidelberg, Brandenburg and Stanley by means of a multiplex PCR (Kim *et al*., 2006). A *S*. Infantis resistant to nitrofurantoin was used for plate count tests.

### Salmonella inoculation in chicken fecal matter

Each *Salmonella* strain was cultured in 4 mL of BHI, then the culture was centrifuged for 5 min at 4,000 x*g*; the supernatant was discarded, and the pellet was re-suspended in 500 μL of sterile saline solution. The process was repeated once to eliminate remnants of culture medium, and the resulting suspension was used to inoculate chicken feces. Fecal material was obtained from 3 2-week old broiler chickens free of *Salmonella*.

Fecal matter from the 3 chickens was pooled and split into seven 10 gram aliquots, placed in seven Petri dishes, mixed with 100 μL of each *Salmonella* strains, and three dishes were left in an aerobic environment and three in anaerobic conditions; all dishes were placed at room temperature and incubated for 24, 48 and 72 hrs (Vasco *et al*., 2015). Anaerobic atmosphere was created by means of BD GasPak™ EZ Anaerobe Gas Generating Pouch System with Indicator.

### Colony counts

Two dishes containing fecal matter with *S.* Infantis (strain POL 398 B) suspension (1.5 × 10^9^ cells per mL) incubated as before under aerobic and anaerobic conditions was subjected to colony count in culture media for *Salmonella*, *E. coli* and coliforms (0 hrs). After incubation, contents of the different dishes were diluted. Dilutions were made up to 10^−8^ in buffered peptone water (BPW), and the dilutions 10^−6^, 10^−7^ and 10^−8^ were inoculated using plate pour method onto XLD and XLD with nitrofurantoin (NIT) (12 mg/L) (Sandegren *et al*., 2008) (we took advantage of the *Salmonella* strain’s resistance to nitrofurantoin to facilitate *Salmonella* colony count). Typical *Salmonella* colonies were counted in XLD and XLD with NIT. The three mentioned dilutions from each one of the fecal matters (aerobic or anaerobic) dishes were inoculated onto 3M™ Petrifilm E. coli/Coliform Count Plates (in duplicate) and incubated for 24 and 48 hrs at 37 °C. We counted red colonies (*E. coli*) from dishes incubated 24 and 48 hrs (from both aerobic and anaerobic). The number of coliforms corresponded to the sum of the red and blue colonies. For each treatment, we included and analyzed a fecal sample without *Salmonella* inoculation as a control.

### Calculation of specific growth rate, μ

The specific growth rate (μ) was calculated using the formula:

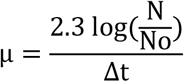

where N is the final population after a time interval of incubation, Δt, and No is the initial population (Maier, 2009; Montville *et al*., 2012).

### Quantitative PCR

We determine the increase in *Salmonella* cells in chicken fecal matter at different time points by the loop mediated isothermal amplification method 3M™ Molecular Detection Assay 2 - Salmonella (MDA2SAL). For this purpose, we performed a calibration curve (Gandelman *et al*., 2010) as follows. We prepared a stool suspension with 1 g of chicken fecal matter and 9 mL of BPW (stool dilution). For the 10^−1^ standard suspension we added to 900 µL of the stool dilution 100 μL of a *Salmonella* Infantis (strain POL 398 B) suspension whose cell concentration was determined by counting in a Petroff-Hausser chamber. For each of the following standard suspensions (10^−2^ to 10^−10^), we used 900 μL of stool dilution and 100 μL of successive dilutions of the *S*. Infantis suspension. Then we mix 20 μL of each standard suspension with the reagents of the kit 3M™ MDA2SAL and analyze them following manufacturer instructions.

Quantitative PCR was carried out after different incubation times (0, 24, 48 and 72 hrs); we made a 10^−1^ dilution of the chicken feces (inoculated with each serovar: Infantis, Dublin, Heidelberg, Brandenburg and Stanley) and analyzed using 3M™ MDA2SAL. We used regression equation to calculate the concentration of *Salmonella* cells at each time point (Gandelman *et al*., 2010). Additionally, for each trial we run suspensions of the strains analyzed, whose concentration (number of cells per mL) was determined in the Petroff-Hausser chamber.

### Statistic analysis

Using SPSS, we performed the U Mann-Whitney test, which compares independent samples that do not have a normal distribution (it is the non-parametric version of the Student’s t-test).

## Results and discussion

We found that *Salmonella* Infantis inoculated in chicken fecal matter multiplied, both in aerobic and anaerobic conditions, however, the aerobic growth was higher, notably from 48 to 72 hrs. Conversely, *E. coli* growth reached its peak after 48 hrs, from then on, the growth decreased (Fig. 1). Coliforms grew similarly but their numbers were higher (Supporting Information Fig. S1).

**Fig. 1.**
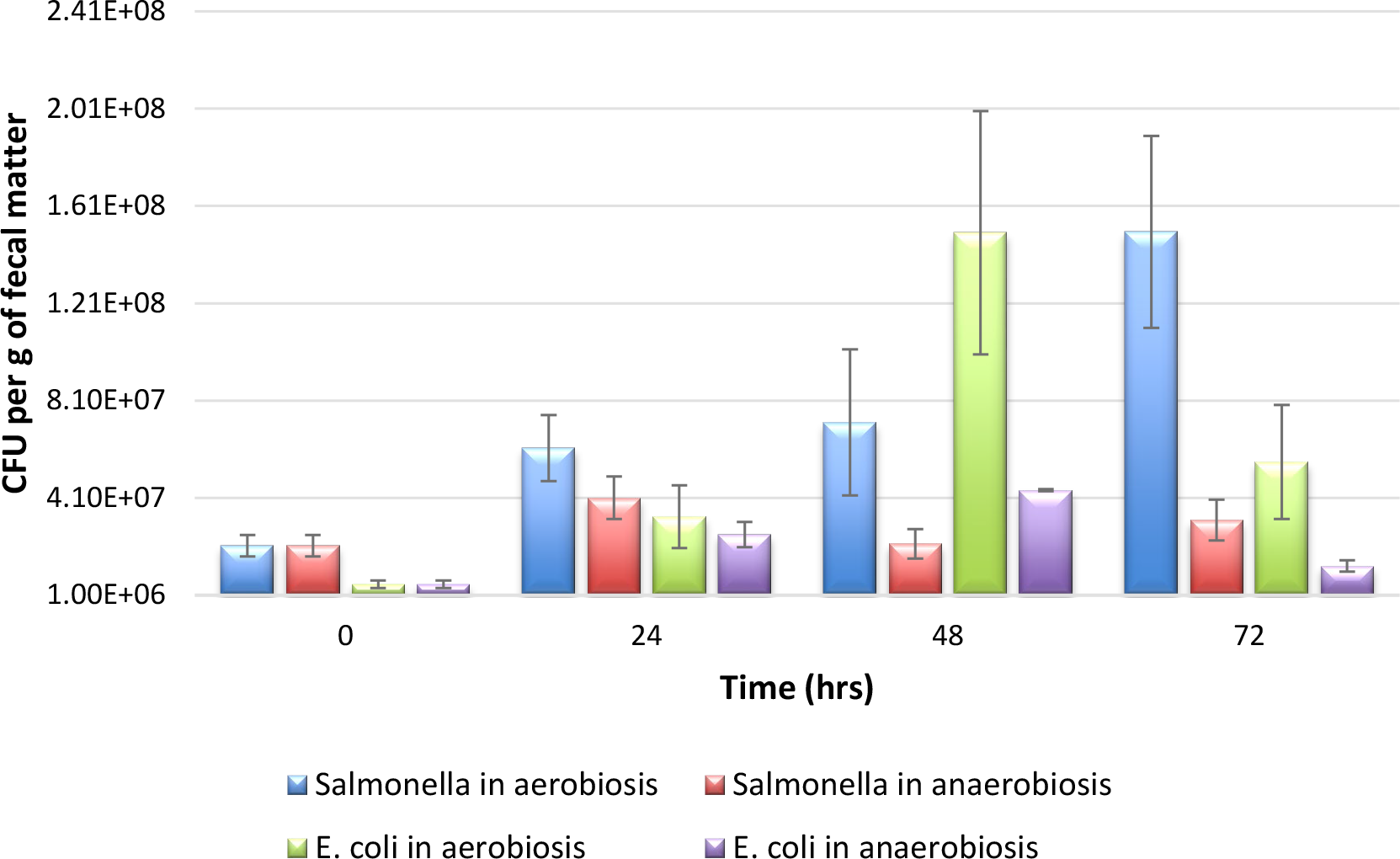
Growth of *Salmonella* Infantis and *E. coli* in chicken fecal matter, under aerobic and anaerobic conditions. Typical *Salmonella* colonies were counted in XLD and XLD with NIT (12 mg/L) (we took advantage of the *Salmonella* strain′s resistance to nitrofurantoin to facilitate *Salmonella* colony count). *E. coli* was counted in 3M™ Petrifilm E. coli/Coliform Count Plates (in duplicate). Bars indicate standard error.

To determine whether this aerobic growth pattern could be extrapolated to other *Salmonella* serovars we inoculated chicken fecal matter with *Salmonella* strains (belonging to 5 different serovars), run the loop mediated isothermal amplification 3M™ Molecular Detection Assay 2 - Salmonella (MDA2SAL) at different times (under aerobiosis and anaerobiosis), and found that all strains showed similar behavior (Fig. 2).

**Fig. 2.**
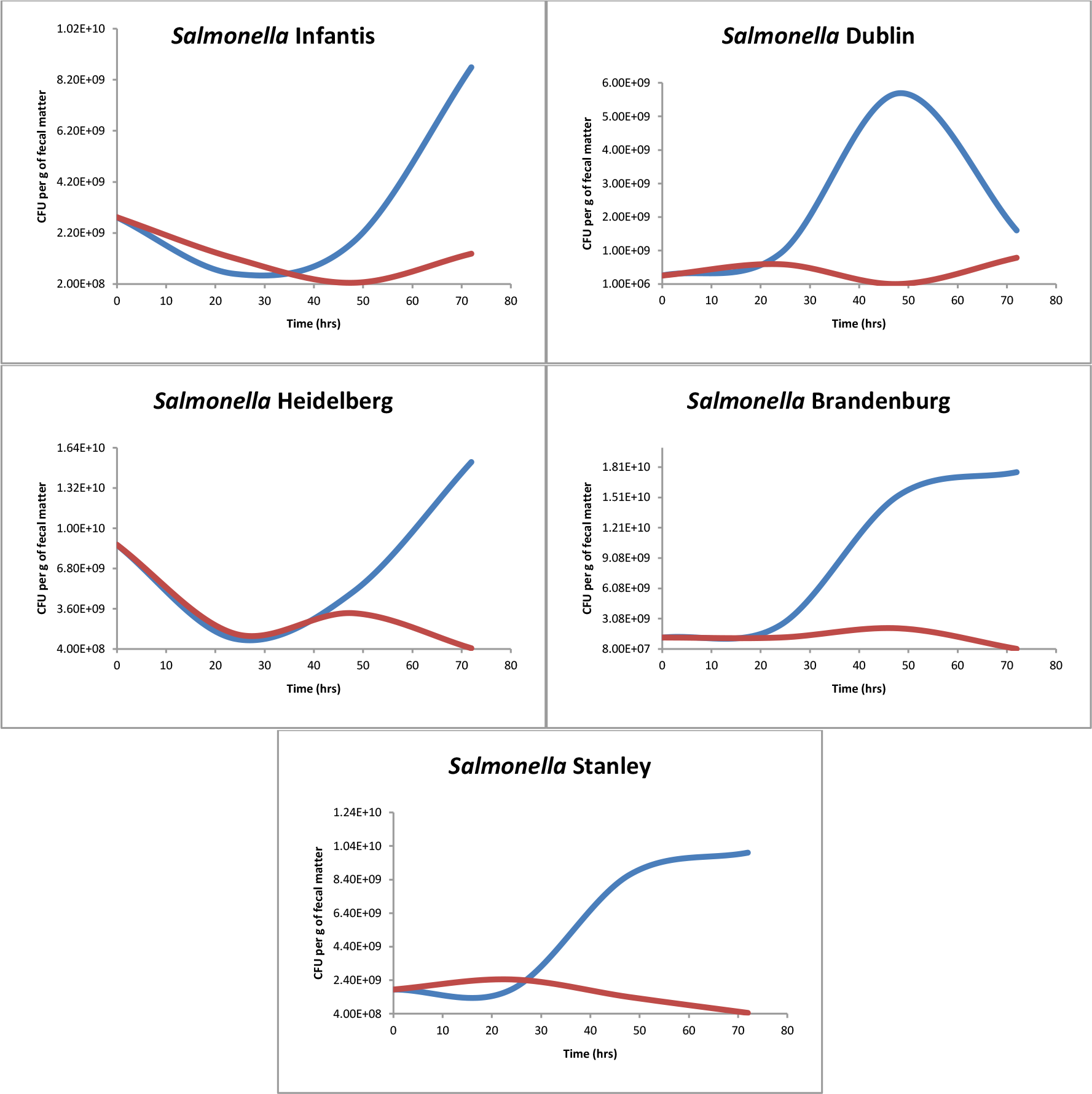
Growth curves of five *Salmonella* serovars. These curves were obtained for five *Salmonella* serovars by 3M™ Molecular Detection Assay 2 - Salmonella (MDA2SAL). The blue lines correspond to the growth under aerobic conditions and red ones, to the growth under anaerobic conditions.

*Salmonella* spp. growth under aerobic conditions continued for 72 hrs (Fig. 3).

**Fig. 3.**
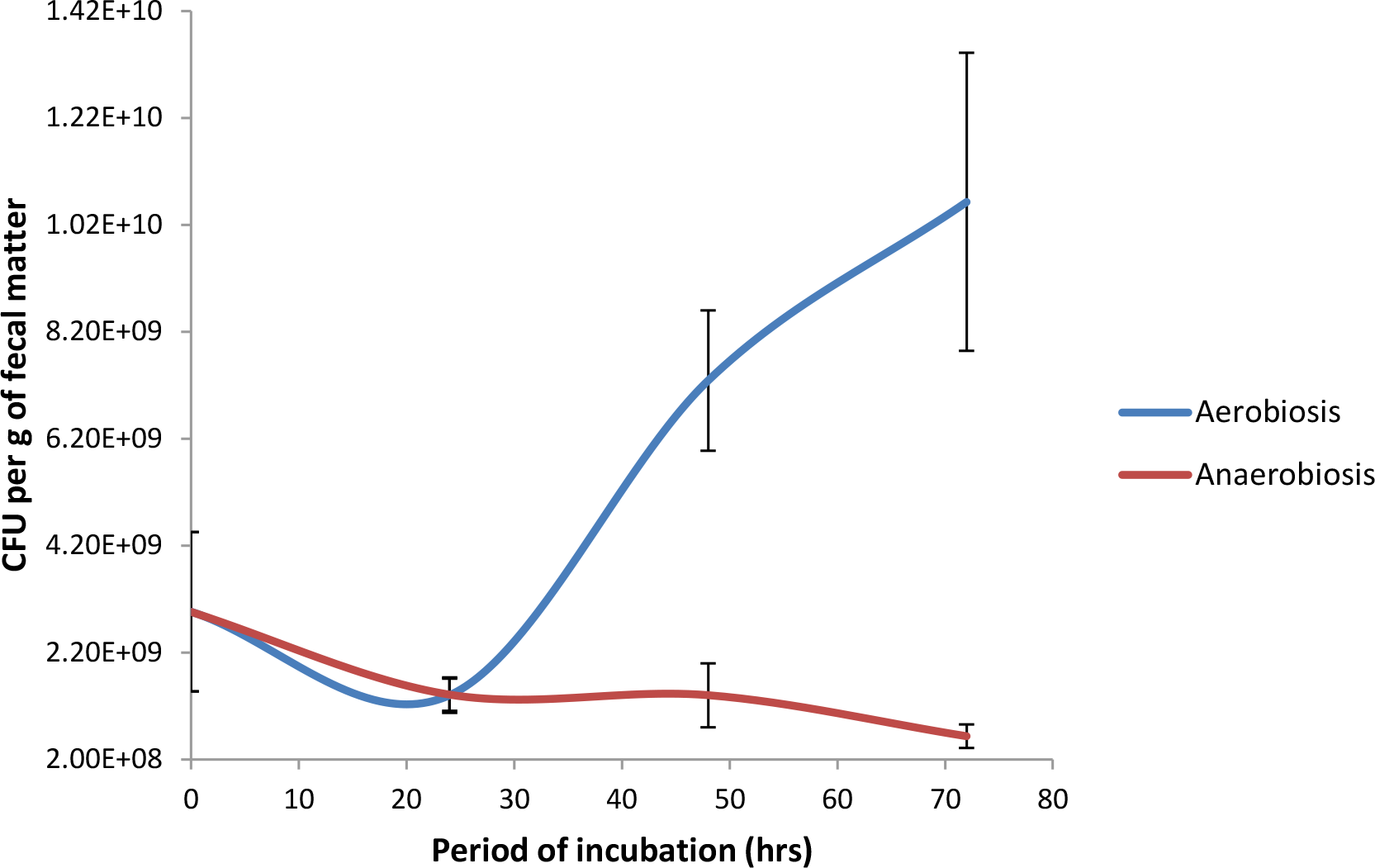
Growth curves of *Salmonella* spp. in chicken fecal matter, under aerobic and anaerobic conditions. Curves were obtained for five *Salmonella* serovars (Infantis, Dublin, Heidelberg, Brandenburg, and Stanley) by 3M™ Molecular Detection Assay 2 - Salmonella (MDA2SAL). Bars indicate standard error.

The specific growth rate (μ) under aerobic conditions showed that during the first 48 hrs *E. coli* had the highest value decreasing during the following 24 hrs, unlike *Salmonella* which started fast growth at this time (Fig. 4). We found that both *Salmonella* and *E. coli* multiplied in fecal matter outside a host. Our results indicate that *Salmonella* (as other *Enterobacteriaceae*) multiplies aerobically in fresh fecal matter (at higher levels than in the intestine) which may be a key step in the infective cycle. Our results show clear evidence that the fecal matter is a transient but very important component of the *Enterobacteriaceae* life cycle, where enterobacterial population expands (Russell and Jarvis, 2001; Vasco *et al.*, 2015; Barrera *et al*., 2018), increasing the chances of reaching other hosts. Our study has an important limitation since we were unable to measure oxygen in fecal matter to confirm our results.

**Fig. 4.**
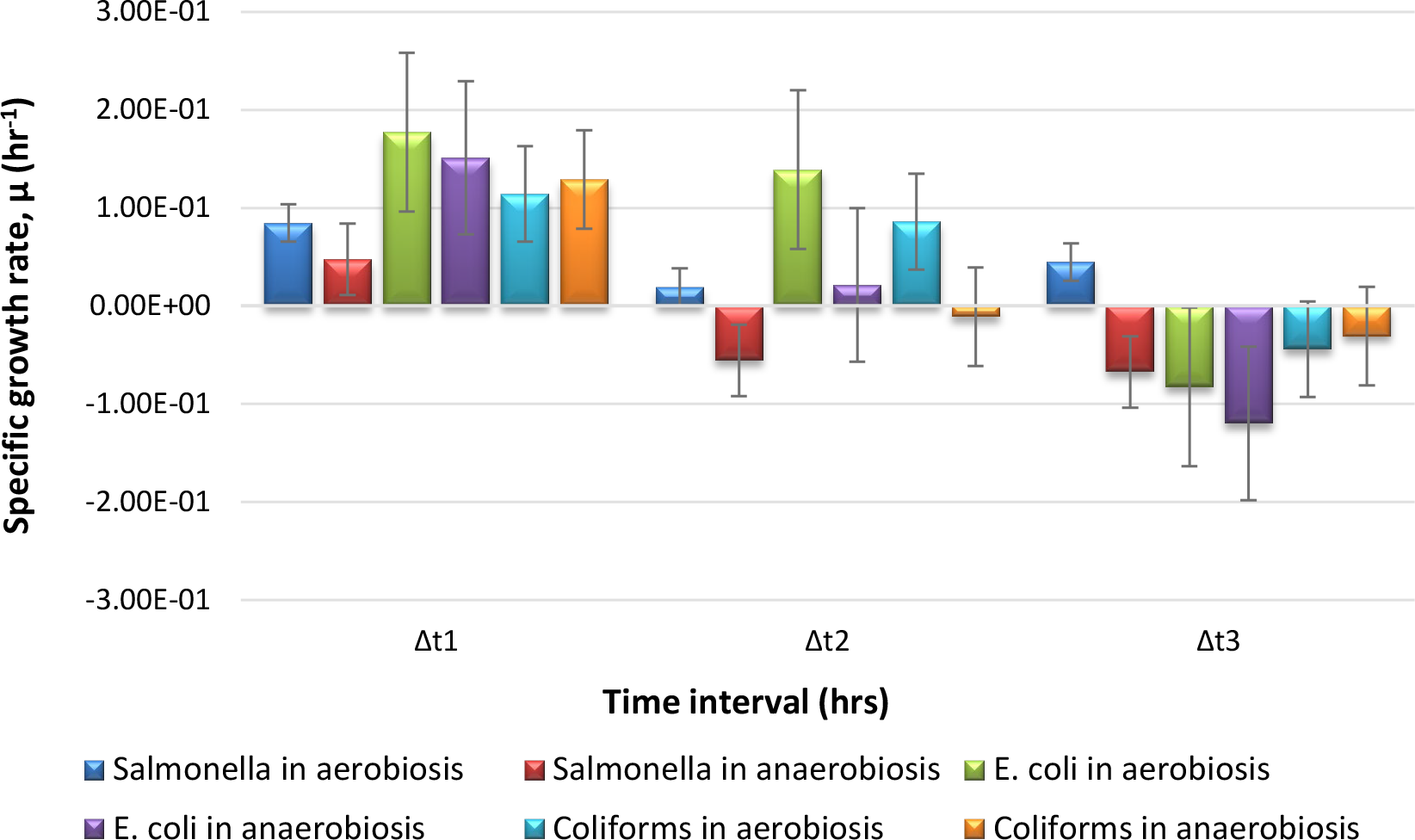
Specific growth rate for *Salmonella*, *E. coli* and coliforms, under aerobic and anaerobic conditions. Specific growth rate, μ, was calculated with the formula: 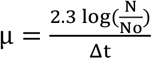, where N is the final population after a time interval of incubation, Δt, and No is the initial population. The incubation times were: t1 = 0 hrs, t2 = 24 hrs, t3 = 48 hrs and t4 = 72 hrs. And the intervals were: Δt1 = t2 − t1, Δt2 = t3 − t2 and Δt3 = t4 − t3. Bars indicate standard error.

Microbiologists have struggled to explain why bacteria adapted to the anaerobic intestinal milieu possess energetically costly machinery to use oxygen (Govantes *et al*., 2000). Further, it has been shown that aerobic respiration is not important for *Salmonella* intestinal colonization (Barrow *et al*., 2015). We hypothesize that the reason for this apparent evolutionary mystery may be related to the enterobacterial ability to grow in fecal matter under aerobic conditions. *Enterobacteriaceae* are facultative anaerobic which can synthesize ATP by different enzymatic pathways, depending on the external concentration of O_2_ and the redox changes in the environment. When O_2_ is available, the bacteria obtain energy by aerobic respiration, with O_2_ being the final acceptor of electrons. In shortage of O_2_, these bacteria generate ATP by one of the following mechanisms: i) synthesis of terminal oxidases that allow the bacteria to take advantage of traces of O_2_; ii) use of other inorganic molecules (such as NO_3_^−^ and S_4_O_6_^2−^) as final electron acceptors (Yamamoto and Droffner, 1985; Bueno *et al*., 2012; Rivera-Chávez *et al.*, 2013); iii) use of organic compounds as donors and acceptors (Madigan *et al*., 2012). Aerobic respiration produces much better performance in terms of ATP molecules per substrate molecule (Madigan *et al*., 2012).

Our results also have critical implications in waste management as we show that aerobic treatments may not be the best to reduce the numbers of *Salmonella* cells (Bueno *et al*., 2012; Rivera-Chávez *et al*., 2013). Additionally, the loose consistency of avian feces allows the entry of air and this phenomenon may contribute to the proficiency of these animals to spread *Salmonella*. Similarly, loose stools, caused by *Salmonella* infection, may favor the growth of this bacterium in fecal matter from animals with different fecal consistency.

Growth curves of *Salmonella* and *E. coli* at different incubation times suggest a negative correlation which may indicate that both species are competing possibly for oxygen, as was described in the intestine (Barrow *et al*., 2015; Velazquez et al., 2019). We speculate that, since *E. coli* multiplies initially more than *Salmonella*, it limits the growth of *Salmonella* until it decreases its growth rate. Once the growth of *E. coli* begins to decrease, *Salmonella* no longer must compete for oxygen and grows faster. Our findings disagree with the notion that once *Salmonella* and *E. coli* are excreted from the host, they enter a viable but not culturable status (Winfield and Groisman, 2003).

## Conflict of interest

The authors declare no conflict of interest.

## Supporting Information

**Fig. S1.**
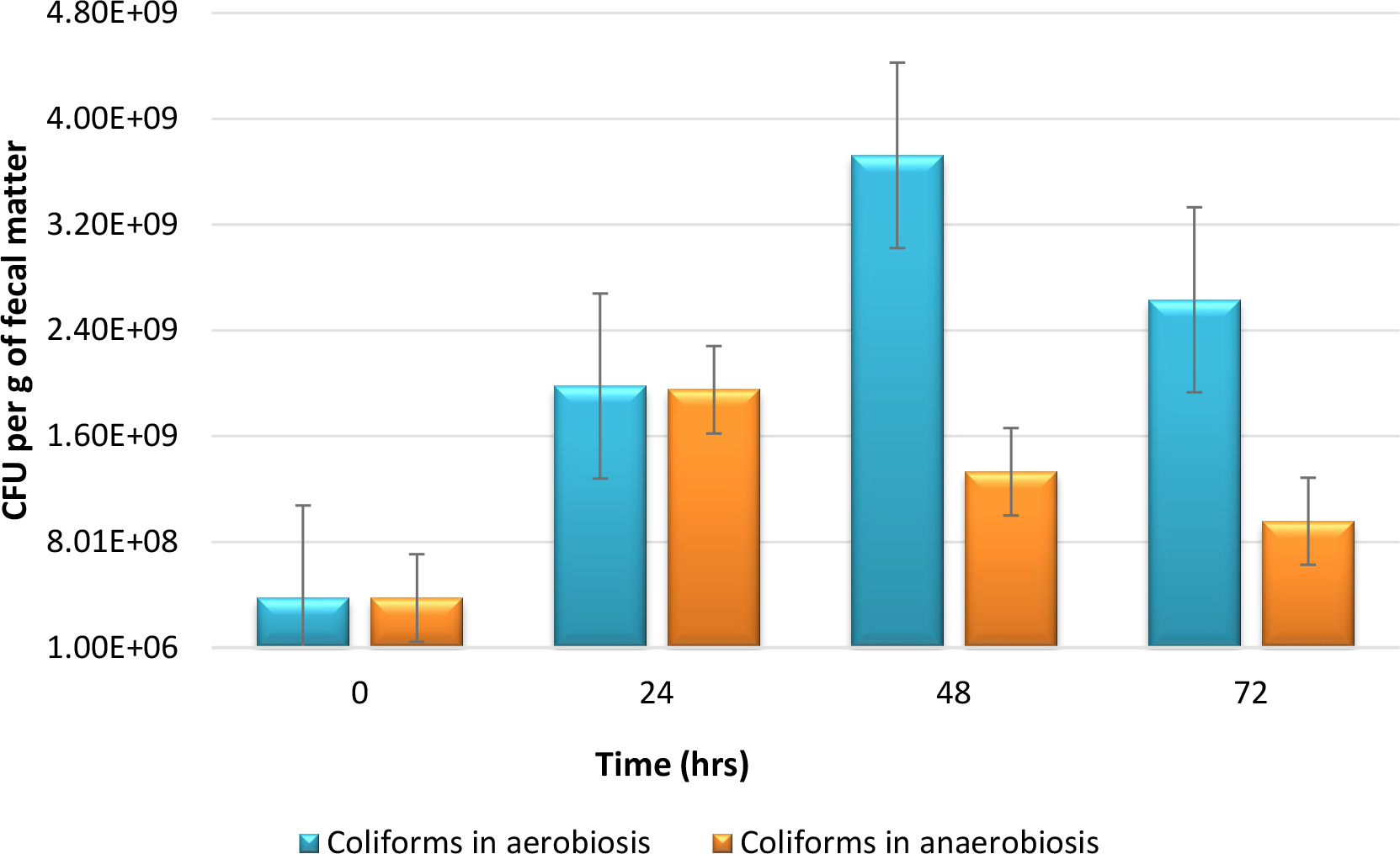
Growth of coliforms in chicken fecal matter, under aerobic and anaerobic conditions. The number of coliforms corresponded to the sum of the red and blue colonies with gas in 3M™ Petrifilm E. coli/Coliform Count Plates incubated 24 and 48 hrs. Bars indicate standard error.

